# Effects of moderate and high intensity isocaloric aerobic training upon microvascular reactivity and myocardial oxidative stress in rats

**DOI:** 10.1101/655530

**Authors:** Lorena Paes, Daniel Lima, Cristiane Matsuura, Maria das Graças de Souza, Fátima Cyrino, Carolina Barbosa, Fernanda Ferrão, Daniel Bottino, Eliete Bouskela, Paulo Farinatti

**Affiliations:** Laboratory for Clinical and Experimental Research on Vascular Biology, Biomedical Center, University of Rio de Janeiro State (UERJ) - **Address**: Rua São Francisco Xavier, Pavilhão Reitor Haroldo Lisboa da Cunha, térreo, sala 524, Maracanã. Rio de Janeiro, RJ, Brasil. CEP: 20550-013; Membrane Transport Laboratory, Department of Pharmacology and Psychobiology, University of Rio de Janeiro State (UERJ) - **Address**: Avenida 28 de Setembro 87, Pavilhão Américo Piquet Carneiro, 5° andar, Vila Isabel, Rio de Janeiro, RJ, Brasil. CEP: 20551-030; Laboratory of Physical Activity and Health Promotion, University of Rio de Janeiro State (UERJ) - **Address**: Rua São Francisco Xavier 524, sala 8121F, Maracanã, Rio de Janeiro, RJ, Brazil, CEP: 20550-900; Graduate Program in Sciences of Physical Activity, Salgado de Oliveira University (UNIVERSO) - **Address**: Rua Marechal Deodoro 211, Centro, Niteroi, RJ, Brasil. CEP: 24030060

**Keywords:** Microcirculation, intravital microscopy, oxidative stress, antioxidant enzymes, energy expenditure

## Abstract

Systemic and central cardiovascular adaptations may vary in response to chronic exercise performed with different intensities and volumes. This study compared the effects of aerobic training with different intensities but equivalent volume upon microvascular reactivity in cremaster muscle and myocardial biomarkers of oxidative stress in Wistar rats. After peak oxygen uptake (VO_2peak_) assessment, rats (n=24) were assigned into three groups: moderate-intensity exercise training (MI); high-intensity exercise training (HI); sedentary control (SC). Treadmill training occurred during 4 weeks, with exercise bouts matched by the energy expenditure (3.0-3.5 Kcal). Microvascular reactivity was assessed *in vivo* by intravital microscopy in cremaster muscle arterioles, while biomarkers of oxidative stress and eNOS expression were quantified at left ventricle and at aorta, respectively. Similar increasing *vs.* sedentary control group (SC) occurred in moderate intensity training group (MI) and high-intensity training group (HI) for endothelium-dependent vasodilation (10^−4^M: MI: 168.7%, HI: 164.6% *vs.* SC: 146.6%, *P*=0.0004). Superoxide dismutase (SOD) (HI: 0.13 U/mg *vs.* MI: 0.09 U/mg and SC: 0.06 U/mg; *P*=0.02), glutathione peroxidase (GPX) (HI: 0.00038 U/mg *vs.* MI: 0.00034 U/mg and SC: 0.00024 U/mg; *P*=0.04), and carbonyl protein content (HI: 0.04 U/mg *vs.* MI: 0.03 U/mg and SC: 0.01 U/mg; *P*=0.003) increased only in HI. No difference across groups was detected for catalase (CAT) (*P*=0.12), Thiobarbituric acid reactive substances (TBARS) (*P* = 0.38) or eNOS expression in aorta (*P*=0.44). In conclusion, higher exercise intensity induced greater improvements in myocardium antioxidant defenses, while gains in microvascular reactivity appeared to rely more on exercise volume than intensity.

## INTRODUCTION

Aerobic training is widely acknowledged as an effective strategy to maintain health and reduce cardiovascular risk factors [1]. Within this context, microvascular endothelial function [2–5] and myocardial antioxidant defenses [6–9] have been extensively investigated, as reflecting early systemic and central cardiovascular changes [2, 6, 10, 11]. In regards to endothelial function, albeit chronic exercise seem to improve vasodilation of microvessels due to shear stress and circulating factors [2, 12], some research suggested that too vigorous training may increase oxidative stress and inflammation, therefore leading to deterioration of endothelial function [13, 14]. On the other hand, myocardium integrity seems be favored by high-intensity training [15], due to greater production of antioxidants protecting against reactive oxygen species (ROS) [7, 16]. In short, different exercise intensities may elicit dissimilar chronic effects upon systemic and central cardiovascular markers – although endothelial function may be jeopardized by high-intensity training [14, 17], myocardium antioxidant protection could be benefited [15].

Particular attention should be therefore given to exercise intensity and volume, for a better understanding of benefits and risks related to those variables. A possible approach to address this question would be to compare the effects of aerobic training performed with different intensities, but similar overall volume as defined by the energy expenditure (EE) – in other words, isocaloric training bouts. Given that improvements in endothelial function may rely on the amount of EE during exercise [17, 18], it is feasible to speculate that isocaloric protocols would be able to induce favorable effects regardless of training intensity [19]. Moreover, this approach would help to avoid bias, since protocols with higher intensities can also be related to greater EE. This confounding factor precludes isolating the specific effects of exercise intensity upon endothelial function or myocardium integrity.

To date no study using animal models investigated the relative effects of exercise intensity and volume upon endothelial function of microvessels (systemic cardiovascular marker), and antioxidant protection in myocardium (central cardiovascular marker). Thus, the present study aimed to investigate the effects of aerobic training performed with different intensities but equivalent volume, on microvascular reactivity in striated muscle and biomarkers of oxidative stress in myocardium of Wistar rats.

## METHODS

### Ethical approval

Twenty-four male Wistar rats (*Rattus norvegicus*, Anilab, RJ, Brazil) were kept under 12:12-hour light-dark cycle in a temperature-controlled environment (22°C) with free access to water and standard rat chow (Nuvital™, Curitiba, PR, Brazil). Experiments were performed according to principles of laboratory animal care (NIH pub. No. 86-23, revised 1996) and the protocol was approved by the Ethical Committee of the University of Rio de Janeiro State (License number: 024/2015).

### Study Design

After assessment of oxygen uptake at rest (VO_2rest_) and maximal exercise (VO_2peak_), animals (270 g, 12 weeks old) were randomly assigned to three groups: (a) moderate-intensity exercise training (MI; n= 8); (b) high-intensity exercise training (HI; n= 8) and (c) sedentary control (SC; n= 8). Two isocaloric exercise bouts were performed after maximal exercise test to match the duration of training sessions according to the overall EE. After this, HI and MI underwent exercise bouts during four weeks on a motorized treadmill. At the end of training period, cardiorespiratory fitness and microvascular reactivity were assessed *in vivo.* eNOS expression was analyzed from aorta fragments. Left ventricle fragments were collected and immediately frozen in liquid nitrogen for measuring biomarkers of oxidative stress (antioxidants and oxidized biomolecules).

### Maximal Graded Exercise Test

Oxygen uptake at rest (VO_2rest_) and during maximal exercise (VO_2peak_) were determined by indirect calorimetry via metabolic cart (Oxylet™, Panlab Harvard Apparatus, Barcelona, Spain). The gas analyzer was coupled to a treadmill in a Plexiglas chamber, connected through a tube to an air pump used to maintain the airflow inside the chamber. The gas analyzer continuously measured relative concentrations of oxygen (O_2_) and carbon dioxide (CO_2_) effluent in the chamber. The VO_2_ was calculated by specific software (Metabolism™, Panlab Harvard Apparatus, Barcelona, Spain) using equations described elsewhere [20]. Standard conditions of temperature, pressure and humidity (*STPD*) were kept in all experiments.

The VO_2_rest was assessed during a 30 min period with only data of the last 5 min being averaged and recorded as result. Prior to VO_2peak_ assessment in maximal exercise testing, rats underwent treadmill adaptation sessions during three days, with speed set at 16 cm/s during 10-15 min. The testing protocol consisted of load increments of 8 cm/s every 3 min, until the rats were no longer able to run. Exhaustion was determined when animals remained at the end of treadmill (electrical shock grid) for 5 seconds and VO_2peak_ corresponded to the highest VO_2_ obtained during the test [20].

### Training Protocol (Isocaloric Exercise Bouts)

The target workload during exercise bouts was calculated using the oxygen uptake reserve (VO_2_R) method, as previously described [19]: VO_2_R = (fraction intensity) (VO_2peak_ – VO_2rest_) + VO_2rest_, where VO_2peak_ corresponded to the highest VO_2_ during maximal exercise testing. The relative intensity was defined according to each group; animals assigned to MI and HI exercised at speeds corresponding to 50% and 80% of VO_2_R, respectively. Running speeds corresponding to relative intensities were individually calculated, based on VO_2_ obtained in exercise testing.

The duration of isocaloric bouts was calculated (predicted) from values of VO_2_R and then converted to EE, as described elsewhere [20]. In order to confirm that both MI and HI bouts elicited similar EE, the VO_2_ was measured throughout two exercise bouts (test and retest). When it was necessary, adjustments in predicted duration were made before the second bout, to ensure that rats would perform isocaloric exercise bouts with different intensities. Since EE equivalence was confirmed in the retest session, HI and MI groups performed isocaloric training sessions on motorized treadmill without indirect calorimetry measurements (Insight Scientific Equipments™, São Paulo, SP, Brazil) during four weeks, five times a week, according to individual values of duration and running speed. In order to assess the excess post-exercise oxygen consumption (EPOC), the VO_2_ was assessed during 30 min following the isocaloric bouts, in a randomized subgroup of four animals selected from HI and MI.

A standardized surgical procedure to evaluate microvascular reactivity in cremaster muscle were performed [21, 22]. In brief, rats were anesthetized with ketamine and xylazine (65 and 10 mg·kg^−1^ respectively), and connective tissue was separated of cremaster. The muscle was exposed on glass stage surface by pins fixed in edges of tissue. The cremaster was continuously superfused at a rate of 4 ml/min by HEPES-supported HCO_3_^−^-buffered saline solution [composition in mM: NaCl 110.0, KCl 4.7, CaCl_2_ 2.0, MgSO_4_ 1.2, NaHCO_3_ 18.0, N-2-hydroxyethylpiperazine-N′-2ethanesulfonic acid (HEPES) 15.39 and HEPES Na^+^-salt 14.61] bubbled with 5% CO_2_–95% N_2_. The pH was set at 7.4 and the temperature of superfusion solution was maintained at 37.5°C. The preparation was placed under an intravital microscope (Leica™ DMLFS, optical magnification ×600, NA 0.65, Wetzlar, Germany) coupled to a closed-circuit TV system, in order to record images of arterioles. The cremaster preparation was maintained during 30 min at rest, before starting experimental protocol to evaluate microvascular reactivity, as described elsewhere [22, 23].

### Microvascular Reactivity Assessment

Three arterioles (2^nd^ and 3^rd^ order) were selected to be analyzed in each cremaster preparation. After 30 min of rest, images of each arteriole were collected at baseline and after topical application of Acetylcholine (ACh) and Sodium Nitroprusside (SNP) (Sigma-Aldrich, St. Louis, MO, USA) at 10^−8^, 10^−6^ and 10^−4^ M. ACh and SNP were used to evaluate endothelium dependent and independent vasodilations, respectively. Each application was performed during 10 min, with a syringe infusion pump (model 55-2222, Harvard Apparatus™, Boston, MA, USA), producing a cumulative dose–response curve. Internal diameter of arterioles in each moment was measured by specific software (Image J ™, U.S. NIH, Bethesda, MD, USA).

### Expression of eNOS (Western Blot Analysis)

Thoracic aorta was dissected and protein extracted, as previously described [23] [24]. Briefly, aorta was lysed in 50 mM HEPES (pH 6.4), 1% Triton X-100, 1 mM MgCl2, 10 mM EDTA, 1 mg/ml DNase, 0.5 mg/ml RNase containing the following protease inhibitors: 1 mM benzamidine, 1 mM PMSF, 1 mM leupeptin, and 1 mM soybean trypsin inhibitor (Sigma-Aldrich, St. Louis, MO, USA). Protein content was measured using Pierce™ BCA Protein Assay Kit (Thermo Fisher Scientific, MA, USA) and samples containing 50 μg of protein were resolved by electrophoresis (7.5% SDS-PAGE), transferred to PVDF membranes and stained with Ponceau to verify whether the same quantity of protein is present in all lanes. Proteins in PVDF membranes were probed with mouse monoclonal anti-eNOS (1:1000; Becton Dickinson, NJ, USA), incubated overnight at 4 °C. A Ponceau Red staining was used as loading control. After extensive washings in TBS-Tween, PVDF membranes were incubated for 2 h at room temperature with horseradish peroxidase-conjugated secondary antibody anti-mouse IgG, diluted 1:5,000 and developed using Amersham ECL Western Blotting Detection Kit system (GE Healthcare Life Sciences, PA, USA).

### Enzymatic assays and oxidative damage

Left ventricular tissue was dissected and homogenized (about 200 mg of tissue) on ice in PBS buffer 0.1 M (0.1 M NaCl, 0.1 M NaH_2_PO_4_.H2O, 0.1 M NaH_2_PO_4_.2H_2_O, 0.1 M KCl, 6 mM EDTA, pH 7.5). Samples were centrifuged at 5,000 rpm for 20 min at 4 °C and supernatant was collected. All samples were stored at −80 °C for further analysis of enzymatic assays and oxidative damage. SOD, GPX and CAT activity were evaluated in left ventricle homogenate. Results are expressed as U/g of protein. Total protein content was quantified using the BCA assay kit (Bioagency™, Sao Paulo, Brazil).

Measurement of SOD activity is based on its inhibition by pyrogallol autoxidation and assessed by spectrophotometric readings at 420 nm during 5 min [24, 25]. Catalase activity was assessed by standard methods, as described elsewhere [26]. Briefly, this method is based on the rate of hydrogen peroxide decomposition, following the decay in absorbance at 240 nm during 1 minute. GPx activity was assessed by the rate of NADPH disappearance, measured by spectrophotometry (340 nm, during 3 min reading) [27].

The oxidative damage of proteins was assessed through formation of carbonyl groups based on the reaction with dinitrophenylhydrazine (DNPH) [28]. Carbonyl contents were determined by spectrophotometry at 370 nm. Lipid membrane damage was quantified by the formation of byproducts of lipid peroxidation (malondialdehyde, MDA), which are thiobarbituric acid reactive substances (TBARS). The MDA reacts with thiobarbituric acid resulting in a pinkish substance, which is subsequently analyzed by spectrophotometry [29]. TBARS were determined by reading the absorbance at 532 nm (Fluostar Omega™, BMG Labtech, Ortenberg, Germany).

### Statistical Analysis

Normal distribution was ratified by the Kolmogorov-Smirnov test for data regarding maximal graded tests and isocaloric exercise bouts, which are presented as mean ± SEM. Microvascular reactivity are presented as median (1^st^ − 3^rd^ quartile). Comparisons between pre *vs.* post-exercise training were performed for VO_2_ and maximal speed by means of 2-way repeated measures ANOVA, followed by LSD post hoc testing in the event of significant *F* ratios. Microvascular reactivity and biomarkers of oxidative stress were compared only at post training, using Kruskal-Wallis test followed by Dunn test as post hoc verification. In all cases significant level was set at *P* ≤ 0.05 and calculations were performed using the Statistica 10.0 software (Statsoft™, Tulsa, OK, USA).

## RESULTS

Table 1 exhibits data of cardiorespiratory fitness, assessed by a maximal graded test before and after training. At baseline, VO_2peak_ (*P* = 0.98) and maximal running speed (*P* = 0.38) were similar across groups. After training, VO_2peak_ decreased in SC (*P* = 0.007), increased in HI (*P* = 0.001) and increased twice in HI than MI, although this difference lacked of statistical significance (VO_2peak_ Δ = 4.9 vs. 2.2; *P* = 0.12). On the other hand, the increase in maximal speed was significantly greater in HI than MI (*P* = 0.016) as well as HI than SC (*P* = 0.02). Despite the differences detected for VO_2_ with training, gains in body mass gain were similar across groups during all experimental period (SC: 276 ± 65; MI: 272 ± 62; HI: 269 ± 61; P = 0.86).

**Table 1.**
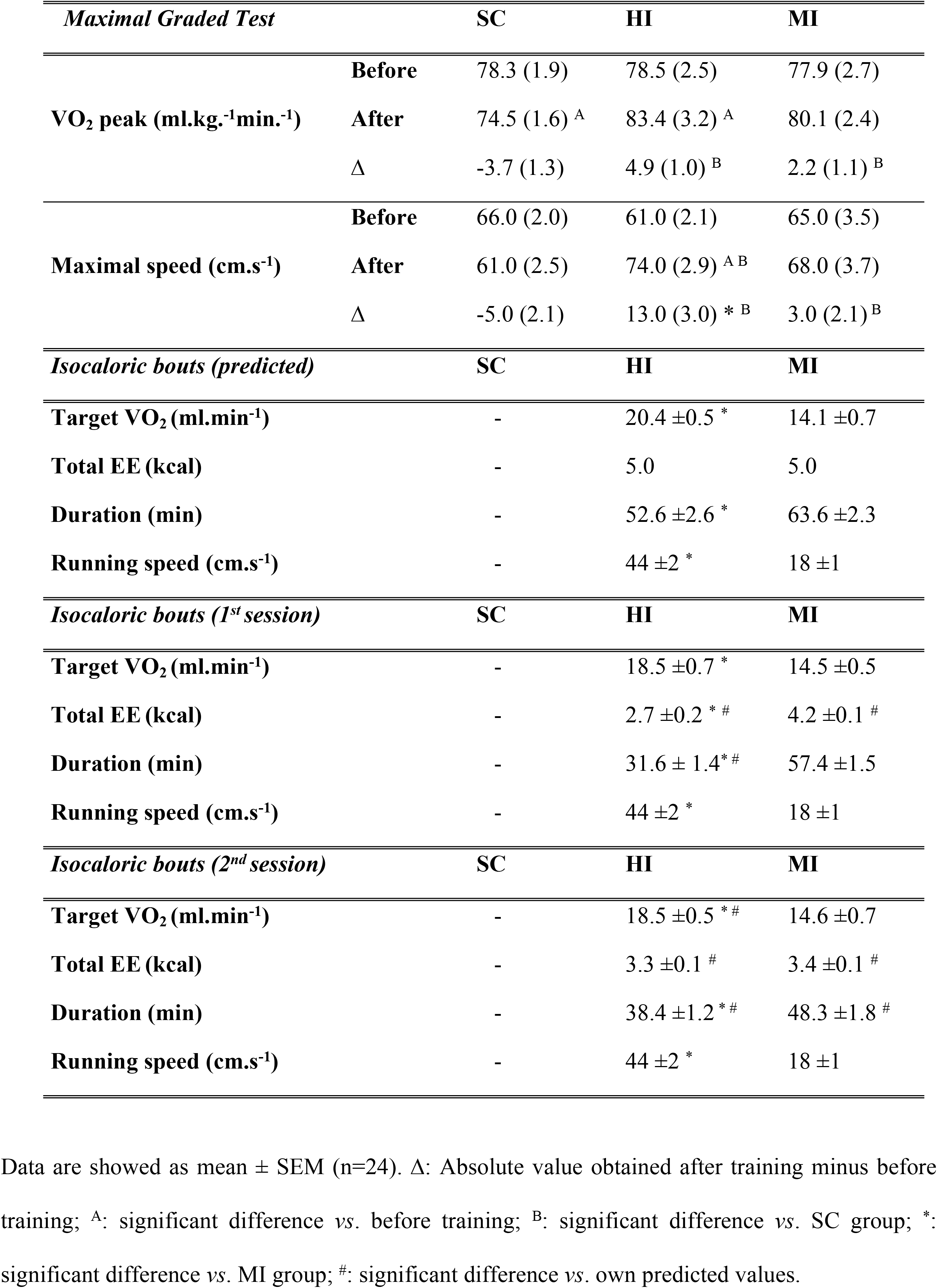
Data from maximal graded exercise test and isocaloric exercise training protocol

| *Maximal Graded Test* |  | SC | HI | MI |
| --- | --- | --- | --- | --- |
| **VO<sub>2</sub> peak (ml.kg.<sup>-1</sup>min.<sup>-1</sup>)** | **Before** | 78.3 (1.9) | 78.5 (2.5) | 77.9 (2.7) |
|  | **After** | 74.5 (1.6) <sup>A</sup> | 83.4 (3.2) <sup>A</sup> | 80.1 (2.4) |
|  | **Δ** | -3.7 (1.3) | 4.9 (1.0) <sup>B</sup> | 2.2 (1.1) <sup>B</sup> |
| **Maximal speed (cm.s<sup>-1</sup>)** | **Before** | 66.0 (2.0) | 61.0 (2.1) | 65.0 (3.5) |
|  | **After** | 61.0 (2.5) | 74.0 (2.9) <sup>A B</sup> | 68.0 (3.7) |
|  | **Δ** | -5.0 (2.1) | 13.0 (3.0) * <sup>B</sup> | 3.0 (2.1) <sup>B</sup> |
| *Isocaloric bouts (predicted)* |  | SC | HI | MI |
| **Target VO<sub>2</sub> (ml.min<sup>-1</sup>)** |  | - | 20.4 ±0.5 * | 14.1 ±0.7 |
| **Total EE (kcal)** |  | - | 5.0 | 5.0 |
| **Duration (min)** |  | - | 52.6 ±2.6 * | 63.6 ±2.3 |
| **Running speed (cm.s<sup>-1</sup>)** |  | - | 44 ±2 * | 18 ±1 |
| *Isocaloric bouts (1<sup>st</sup> session)* |  | SC | HI | MI |
| **Target VO<sub>2</sub> (ml.min<sup>-1</sup>)** |  | - | 18.5 ±0.7 * | 14.5 ±0.5 |
| **Total EE (kcal)** |  | - | 2.7 ±0.2 * # | 4.2 ±0.1 # |
| **Duration (min)** |  | - | 31.6 ± 1.4* # | 57.4 ±1.5 |
| **Running speed (cm.s<sup>-1</sup>)** |  | - | 44 ±2 * | 18 ±1 |
| *Isocaloric bouts (2<sup>nd</sup> session)* |  | SC | HI | MI |
| **Target VO<sub>2</sub> (ml.min<sup>-1</sup>)** |  | - | 18.5 ±0.5 * # | 14.6 ±0.7 |
| **Total EE (kcal)** |  | - | 3.3 ±0.1 # | 3.4 ±0.1 # |
| **Duration (min)** |  | - | 38.4 ±1.2 * # | 48.3 ±1.8 # |
| **Running speed (cm.s<sup>-1</sup>)** |  | - | 44 ±2 * | 18 ±1 |

Data are showed as mean ± SEM (n=24). Δ: Absolute value obtained after training minus before training; <sup>A</sup>: significant difference vs. before training; <sup>B</sup>: significant difference vs. SC group; \*: significant difference vs. MI group; #: significant difference vs. own predicted values.

Table 1 also depicts data extracted from the first and second isocaloric bouts (test and retest sessions). As expected, running speeds (*P* = 0.013) and therefore target VO_2_ (*P* = 0.0002) were always higher in HI *vs*. MI. In the first exercise bout, total EE was significantly different (*P* = 0.007) and could not be matched between groups, because animals in HI were not able to complete the predicted exercise duration before exhaustion. The second bout was performed after adjustments (by reducing duration for MI group) and differences in total EE between groups were no longer detected (*P* = 0.61), albeit target VO_2_ in HI has remained higher than MI (*P* = 0.007). This is reinforced by the EPOC, which was significantly influenced by exercise intensity. At the beginning of recovery, HI had significantly higher VO_2_ than MI (HI = 17.58 ± 1.4 mL/min vs. MI = 13.15 ± 0.8 mL/min vs.; *P* = 0.03) and this pattern was extended until the end of recovery, as shown by the VO_2_ range (HI = 7.85 ± 0.6 mL/min vs. MI = 4.45 ± 0.8 mL/min; *P* = 0.01).

Changes in arterioles diameter in relation to basal conditions (considered as 100%) under topical application of ACh and SNP are depicted in Figure 1. Baseline refers to a period before ACh or SNP application and after 30-min accommodation; at this phase, mean diameter was always similar across groups (SC: 71.73 ± 2.9 μm, MI: 74.91 ± 3.0 μm, HI: 70.55 ± 4.9 μm; *P* = 0.54). Endothelium-dependent vasodilation in response to the highest ACh concentration (10-4M) and to the intermediate concentration (10^−6^M) was significant lower in SC than MI (*P*=0.0016 and *P* = 0.003) and HI (*P* = 0.0012 and *P* = 0.01). In response to the lowest concentration of ACh (10^−8^M), only HI presented significant increased vasodilation in comparison to SC (*P* = 0.02). No significant differences were observed in relation to MI (*P* = 0.17). No difference among groups was detected regarding endothelium-independent vasodilation induced by SNP at any concentrations (10^−8^M: *P* = 0.34; 10^−6^M: *P* = 0.49; 10^−4^: *P* = 0.28).

**Fig. 1.**
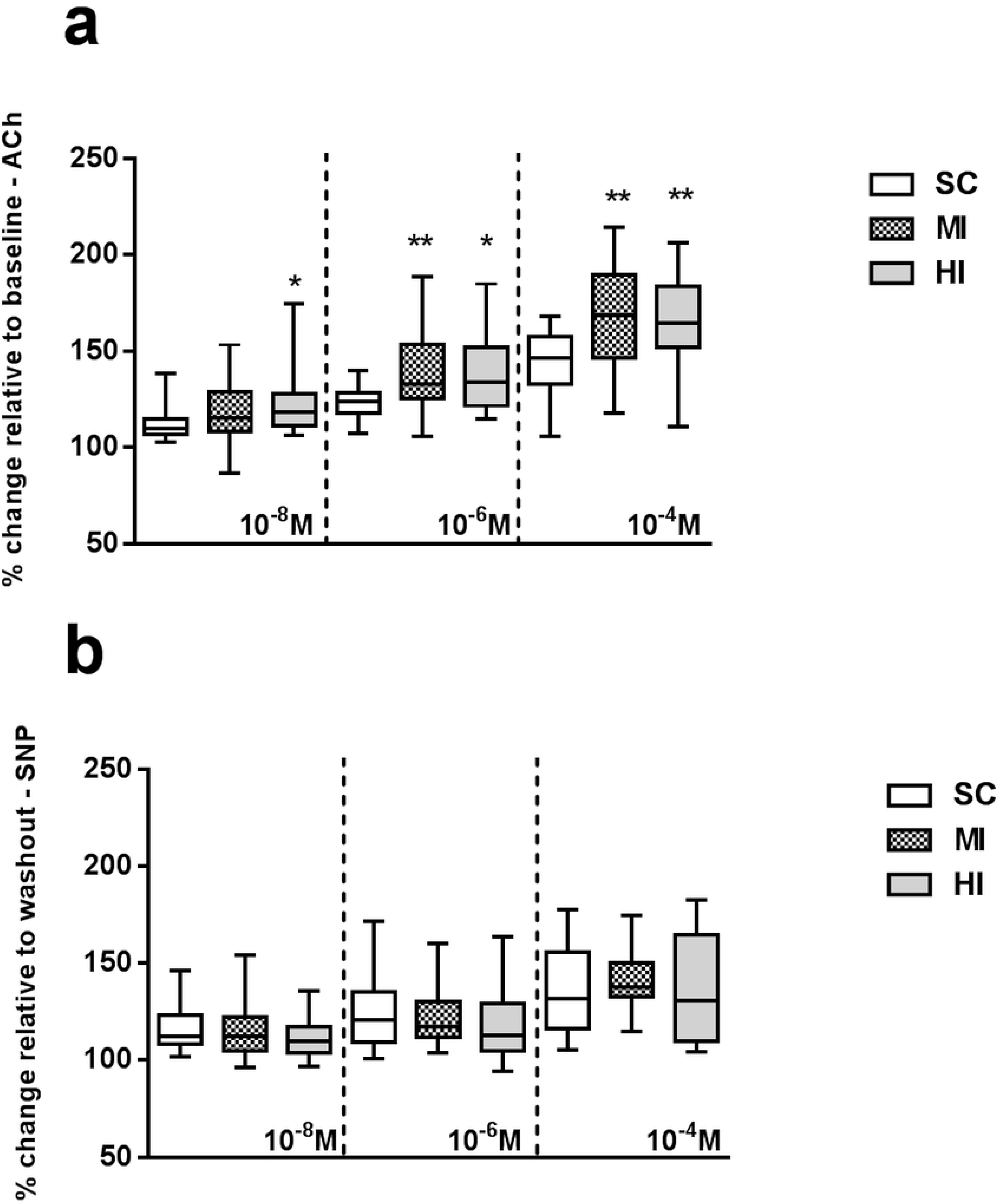
Microvascular reactivity in vivo of arterioles in cremaster muscle. Data are expressed as median (1^st^− 3^rd^quartile) (n = 24). (a) Endothelium-dependent vasodilation; (b) endothelium-independent vasodilation. *Significant difference vs. SC (P < 0.05).

Exercise performed at different intensities elicited different adaptations in antioxidant enzymatic activity and oxidative damage at left ventricle homogenates, as shown in Figure 2. CAT activity did not change in any group (*P* = 0.12). In contrast, SOD and GPX activity were significantly increased in HI (*P* = 0.04; *P* = 0.01), but not MI (*P* = 0.29; *P* = 0.26). Protein carbonyl content was higher in HI (*P* = 0.003), suggesting an increase in protein oxidative damage, whereas no change was observed after training in MI (*P* = 0.07). Malondialdehyde content did not differ among groups (*P* = 0.38). Figure 3 exhibits results of eNOS expression in aorta. Western Blot analysis could not detect differences between groups (HI = 1.10 ± 0.2, MI = 1.39 ± 0.2, SC = 1.00 ± 0.1; P = 0.44).

**Fig. 2.**
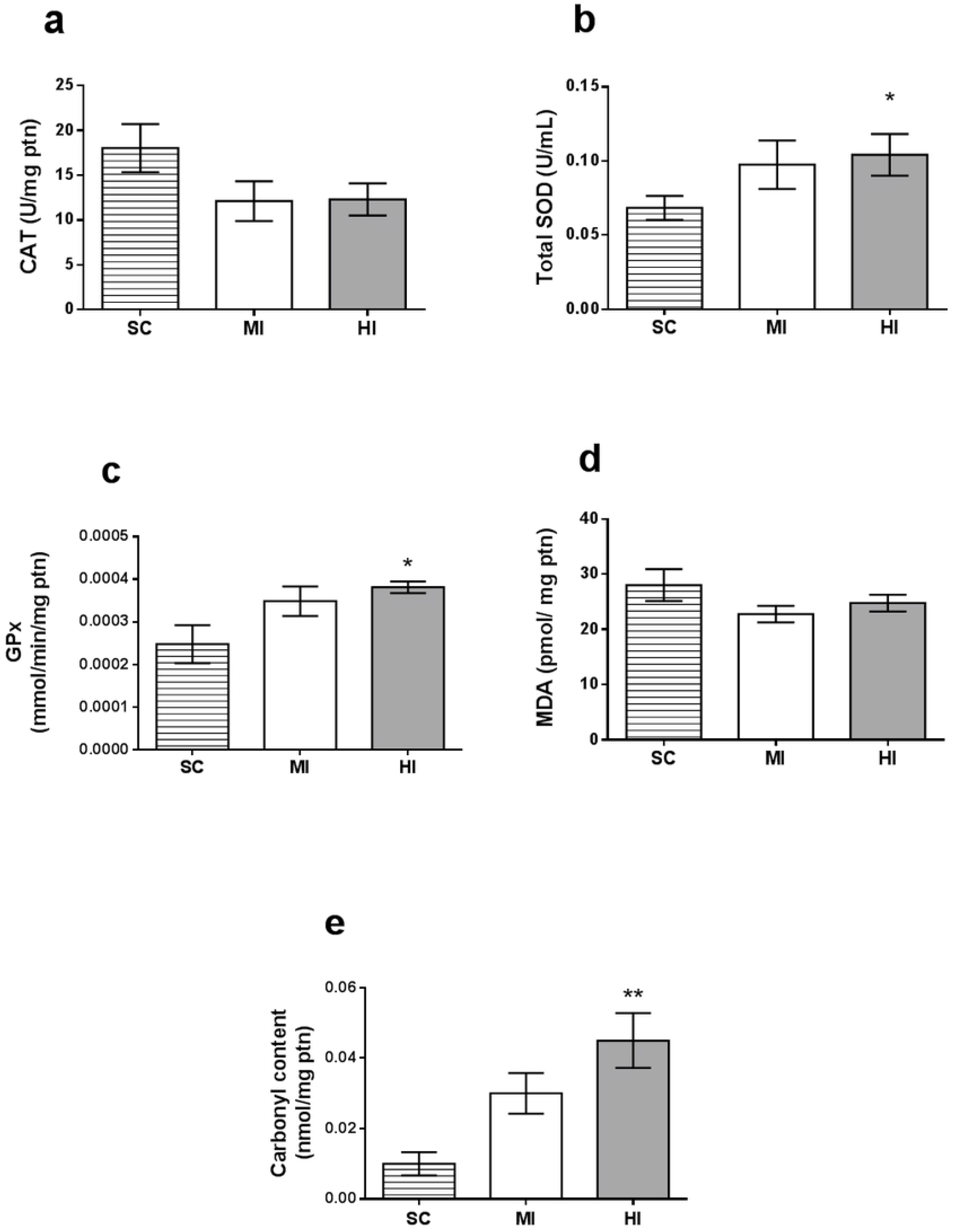
Biomarkers of oxidative stress. Data are expressed as mean ± SEM (n = 24). (a) CAT Activity; (b) SOD Activity; (c) GPX Activity; (d) TBARS - MDA content; (e) Protein Carbonyls. *: significant difference vs. SC group (P < 0.05); **: significant difference vs. SC group (P < 0.01).

**Fig. 3.**
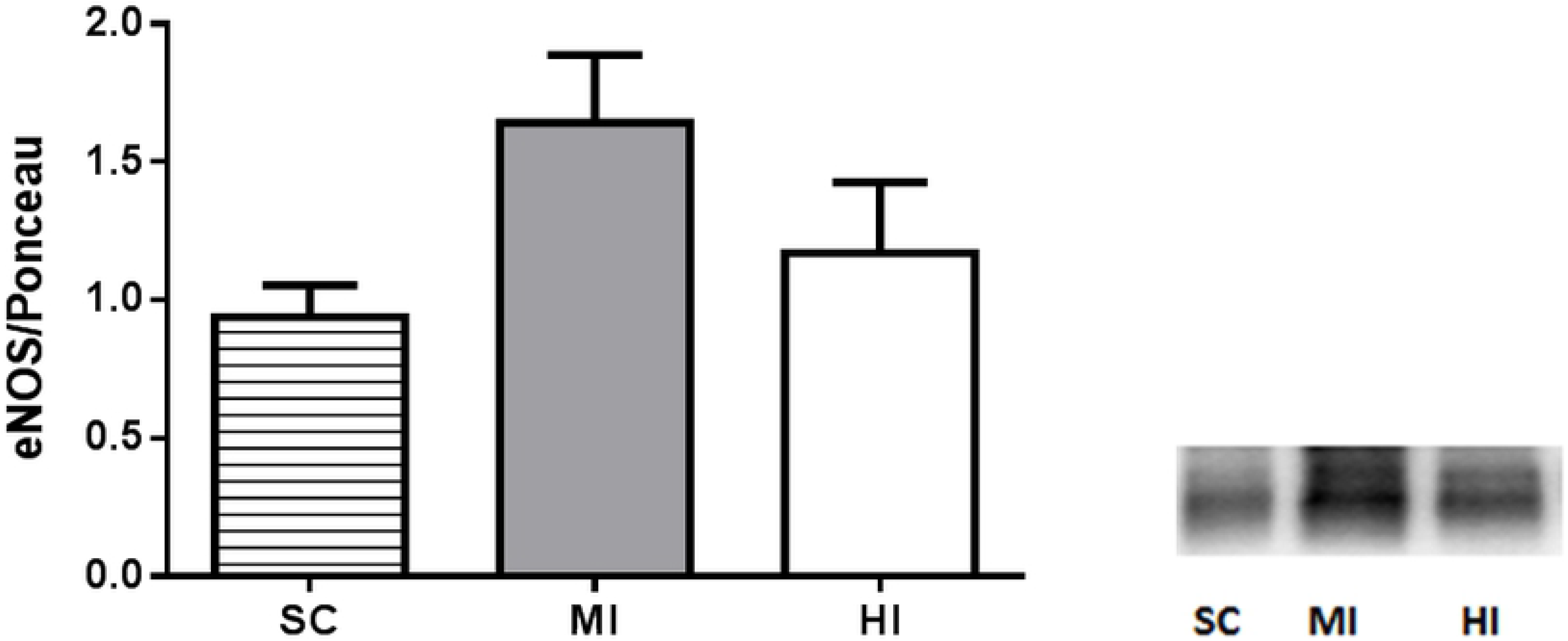
Western blot analysis of aortic eNOS. Data are expressed as means ± SEM (n = 24). No significant difference was found between the groups (P = 0.44).

## DISCUSSION

This study investigated the effects of high- and moderate-intensity aerobic training with equivalent volume (or ‘dose’ reflected by EE) upon microvascular reactivity in cremaster muscle and biomarkers of oxidative stress in myocardium, considered as central and systemic cardiovascular health markers, respectively. It has been hypothesized that protocols with equivalent training volume would elicit similar outcomes, regardless of differences in training intensity. At least for microcirculation our findings confirmed this hypothesis, since improvements in endothelium-dependent vasodilation were similar across training groups. On the other hand, only high-intensity training was able to elicit improvements in SOD, GPX, and protein carbonyl content in myocardium.

Some prior research investigated the effects of different exercise intensities upon isolate vasculature [2, 13, 14] or myocardium [9, 15]. However, we could not find studies assessing the concomitant effects of aerobic training upon those peripheral and central markers. Evidently, this approach seems to be more adequate to investigate whether adaptations induced by different training protocols upon a given marker would extend to others. Moreover, bias related to exercise volume has never been addressed by previous research about effects of exercise intensity upon different cardiovascular markers [14, 30, 31].

Consistently with our initial hypothesis, significant improvements in endothelium-dependent vasodilation were similar across training groups in the highest concentration of ACh (MI = 168.7 *vs.* HI = 164.6; *P* = 0.91), suggesting that effects upon microcirculation would be more related to exercise volume than intensity. This finding concurs with data previously reported, suggesting that potential effects of chronic exercise on vasculature would be rather associated to overall exercise ‘dose’ (as reflected by EE) than isolate relative intensity or duration [17, 18]. Potential mechanisms underlying chronic adaptations in endothelium include increased shear stress and NO bioavailability [2, 32]. Another finding that reinforces the beneficial role of these mechanisms on endothelium function is the lack of differences in endothelium-independent vasodilation between control and trained groups, also in the highest concentration of SNP (SC = 136.1, MI = 141.6, HI = 136.1; *P* = 0.28), which by the way is supported by previous studies [33, 34].

Altogether, those findings support the hypothesis that the caloric cost of physical training *per se* may elicit favorable adaptations in endothelial function. Conversely, our findings did not confirm the premise that microvascular endothelial improvements would be due to greater systemic NO production, since no difference between groups was found for aortic eNOS expression. Several studies indicated that an increase in vascular eNOS would occur in response to prolonged training (>10 weeks) [2]. In the present study, the training duration of 4 weeks was perhaps too short to elicit significant changes in eNOS expression [35].

Some prior research suggested that aerobic training may improve vasoreactivity in arterioles, regardless of concomitant increasing in metabolic activity or blood flow during exercise (i.e. non-exercised muscles) [4, 32, 36]. By choosing the cremaster muscle to assess *in vivo* vasoreactivity instead of active muscle beds (soleus, gastrocnemius) as performed in most studies [2, 32, 34], we reinforced the premise that chronic exercise effects upon microcirculation are rather systemic than local [32, 33]. This systemic response seems to be independent of exercise intensity, which gives room for interesting practical applications in regards to aerobic training and cardiovascular health.

Moreover, our findings suggest that favorable microcirculation adaptations to physical training might occur irrespective of predominance of type I fibers in a given muscle; indeed, slow oxidative fibers are often associated with higher vasodilation capacity [37], while the predominance in cremaster muscle is of fast glycolytic fibers [38]. Despite the overall strengths by deciding for cremaster muscle, the assessment of vasoreactivity solely in such muscle represents the main limitation of the present study.

With respect to the effects of training intensity upon myocardium, only high-intensity training improved SOD and GPx activities, while CAT did not respond to any exercise intervention (*P* = 0.12). These findings were expected and concur with prior studies showing that activity of GPx [9, 15] and SOD [9] increased after vigorous aerobic training, while effects upon CAT remain unclear [15, 39]. Furthermore, the increase of protein carbonyls levels in HI evokes persistent doubts in regards to whether augmentation in tissue damage by ROS-generation in the heart would be necessary to stimulate antioxidant protection, or whether this acute effect would be harmful due to constant cell-damaging [13, 14]. While this issue remains unclear, it is well elucidated that SOD plays an important protective role through superoxide dismutation [15] and GPX is capable of reducing a broad range of hydroperoxides, thereby conferring great protection against cell-damage by oxidation [7, 15].

Some limitations of this study must be mentioned. Firstly, antioxidant activity, lipid peroxidation and protein damage in vasculature have not been assessed. Such biomarkers could indicate whether exercise volume and intensity might also modulate oxidative stress in vessels. Additionally, vasoreactivity was only assessed in the cremaster muscle. Although cremaster is widely used for evaluating systemic vasodilation *in vivo*, its utilization limits generalization and further studies are warranted to investigate the effects of training intensity upon microcirculation in other body regions.

## CONCLUSION

Overall, our findings corroborate the premise that a minimal daily caloric expenditure promoted by exercise would be enough to maintain endothelial health, but not to increase antioxidants in myocardium. Given that gains in antioxidant enzymes were only detected in the group that performed high-intensity training, it seems that vigorous exercise would be necessary to improve antioxidant defenses in myocardium.

## ABBREVIATIONS

eNOS: endothelial nitric oxide synthase
SC: sedentary control group
MI: moderate-intensity exercise training group
HI: high-intensity training exercise group
SOD: superoxide dismutase enzyme
CAT: catalase enzyme
GPX: glutathione peroxidase enzyme
TBARS: thiobarbituric acid reactive substances
ROS: reactive oxygen species
EE: energy expenditure
VO_2_: oxygen uptake
VO_2_R: oxygen uptake reserve
EPOC: excess post-exercise oxygen consumption
ACh: acetylcholine
SNP: sodium nitroprusside
MDA: Malondialdehyde
DNPH: dinitrophenylhydrazine
SEM: standard error of mean.

## AUTHORS’ CONTRIBUTIONS

LP collected, analyzed, and interpreted all data and was a major contributor in writing the manuscript. DL performed the enzymatic assays and oxidative damage and analyzed related data. CM analyzed and interpreted data enzymatic assays and oxidative damage and assisted in editing the manuscript. MDGS assisted in interpretation of data and editing the manuscript. FC assisted in microcirculation experiments and in interpretation of related data. CB performed Western Blot experiments, analyzed and interpreted related data. FF performed Western Blot experiments. DB supervised the overall research project and assisted in editing the manuscript. EB supervised the overall research project and assisted in editing the manuscript. PF supervised the overall project, analyzed and interpreted the data, and was a major contributor in writing the manuscript. All authors read and approved the final manuscript.

## ACKNOWLEDGEMENTS

Authors would like to thank Paulo José Lopes and Claudio Natalino Ribeiro for animal care; João Paulo Barbosa and Lara Serrano for assistance with the animals training.

